# Connexins are essential for the contribution of latent progenitors to healing after spinal cord injury

**DOI:** 10.64898/2026.01.16.699895

**Authors:** María Victoria Falco, Gabriela Fabbiani, Daniel Prieto, Mateo Vidal, Milagros Benítez, María Inés Rehermann, María Constanza Silvera, Renata Simeone, Federico F. Trigo, Raúl E. Russo

## Abstract

Gap junctions are important regulators of the biology of neural stem cells. Both in vertebrates with regenerative abilities and neonatal rodents, ependymal cells communicate via connexin (Cx) 43 and Cx26. Gap junction coupling and Cx26 are down-regulated as the ependyma becomes quiescent in adulthood, but injury overrules this developmental down-regulation suggesting a role for Cx signaling in the reactivation of ependymal cells. However, direct evidence of the involvement of these Cxs in the reactivation of the ependymal stem cell niche is lacking. Here, we aim to explore the role of Cx26 and Cx43 in the response of ependymal cells to injury by means of specific genetic deletion. We find that Cx26 is critical for the proliferative response of ependymal cells to injury. The reduced response of ependymal cells by Cx26 deletion increased the extension of the lesion because of impaired healing resulting from a weak contribution of the ependymal stem cell niche. On the other hand, Cx43 -via gap junction coupling and hemichannels-plays a key role in the communication of Ca^2+^ signals between ependymal cells induced by activation of P2X7 receptors. Our data show that Cxs are relevant targets to manipulate the ependymal stem cell niche to achieve better healing after spinal cord injury.

**Significance statement:** Ependymal cells of the spinal cord are latent progenitors whose manipulation is regarded as a potential strategy to improve healing after injury. Therefore, a better understanding of the mechanisms that reactivate these dormant progenitor-like cells is fundamental. Connexins (Cxs) are key regulators of the biology of neural stem cells and change in the ependyma after injury, suggesting a role in the awakening of the niche. We find here that Cx26 is critical for the proliferative response of ependymal cells and scar formation. Cx43 plays a key role in the early signaling of injury needed to trigger the expression of Cx26. Our work demonstrates that Cxs are relevant targets to manipulate the response of ependymal cells to injury.

## Introduction

The traumatic injury of the spinal cord (SC) leads to devastating conditions because of the disruption of axonal pathways, neuronal death, axonal demyelination and scarring (1). Although mammals lack the ability for regeneration displayed by some non-mammalian vertebrates (2), some cells in the ependyma still react to injury generating new cells that migrate towards the lesion to become astrocytes within the core of the scar (3, 4). The manipulation of ependymal cells is regarded as a promising strategy to potentiate healing (5). To unleash the regenerative potential of the ependymal stem cell niche, a better understanding of the mechanisms that reactivate these dormant progenitor-like cells after injury is needed.

Gap junction coupling has an important role in the regulation of neural stem cells (6), with connexin (Cx) 43 and Cx26 expressed in radial glia during cortical development (7, 8). The role of Cxs as regulators of proliferation is maintained in adult neurogenic niches (9). Both in vertebrates with regenerative abilities and in neonatal rodents, ependymal cells in the SC communicate via Cx43 and Cx26 (10, 11). The incidence of gap junction coupling is developmentally down-regulated as the ependyma becomes quiescent in adulthood, an event paralleled by downregulation of Cx26 (12). At the time injury-induced proliferation peaks (3), gap junction coupling between adult ependymal cells increases, resembling the functional state of the immature ependyma (12). Interestingly, injury overrules the developmental down-regulation of Cx26 in ependymal cells suggesting a role in the awakening of these latent progenitors.

Cx signaling is important for wound healing and scarring in different types of tissues (13). The idea that Cxs play a similar function in the ependymal stem cell niche to regulate endogenous repair is supported by the fact that pharmacological blockade of gap junctions reduced injury-induced proliferation of ependymal cells (12). In epithelia, the wound-induced changes of Cx26 and Cx43 expression occur in a complex spatiotemporal manner (14) and the blockade of Cx43 speeds up wound closure (15). We speculated that Cx26 and Cx43 may play a similar role in healing mediated by ependymal cells. Indeed, a previous study shows that local application of the connexin blocker Gap26 reduces the injury-induced proliferation of ependymal cells (12). However, due to the low selectivity of Gap26 for different Cx isoforms and the fact that pharmacological blockade interferes with Cx hemichannels and gap junctions in different types of cells, the functional relevance of Cx26 and Cx43 in ependymal cells remains unresolved. Here, we aimed to explore the role of Cx26 and Cx43 in the response of the ependyma to injury by means of genetic deletion of specific Cxs in ependymal cells. By applying a multi-technical approach combining both in vivo and in vitro models, we find that injury-induced expression of Cx26 is critical for the proliferative response of ependymal cells to injury, and that its absence generates a severe impairment of scar formation. On the other hand, Cx43 participates in the early cellular signaling of injury and its deletion also reduces the response of the ependyma to traumatic injury. Our data suggests that Cx43 -together with P2X7 receptors-plays a key role in the communication between ependymal cells via Ca^2+^ waves that trigger downstream events such as the Cx26 up-regulation needed for the resumption of proliferation. Collectively, our findings show that both Cx43 and Cx26 are essential elements of the molecular pathways responsible for the reactivation of dormant progenitor-like cells and represent relevant targets that may modulate the response of the ependymal stem cell niche to promote a better repair after spinal cord injury (SCI).

## Results

Gap junction coupling and Cx26 expression decline during post-natal development as the ependymal stem cell niche becomes quiescent in adulthood (12). SCI induces a strong reactivation of progenitor-like cells in the ependyma that is paralleled by the re-expression of Cx26 and an increased incidence of gap junction coupled cells, suggesting Cx26 may be implicated in the mechanisms involved in re-entering the cell cycle. Indeed, the pharmacological blockade of gap junctions and Cx hemichannels with the peptide Gap26 strongly reduces the injury-induced proliferation of ependymal cells (12). However, this pharmacological approach has the limitation that Gap26 blocks all types of Cxs in different types of cells within the SC. To tackle the role of specific Cx isoforms in the response of ependymal cells to SCI, we used a Cre-Lox system to genetically delete Cx26 selectively in ependymal cells by crossing FoxJ1CreER-tdT -which expresses the Cre recombinase under the control of the FoxJ1 promoter active in cells with motile cilia (4,16) - with mice having the Cx26 gene (Gjb2) between LoxP sites (Fig. 1 A). We induced recombination in FoxJ1CreER-tdT-Cx26^fl/fl^ mice by injection of tamoxifen during 5 days and allowed 10 days for tamoxifen clearance before performing a dorsal hemisection of the SC (Fig. 1 B, Fig. S1). The basic characteristics (e.g., expression of Sox2, very low rate of mitotic activity) of the ependyma in the uninjured spinal cord of Cx26^−/−^ mice were essentially similar to those of control animals (Fig. S2 A-B, D, E-F and H). As described before (12), a dorsal hemisection induced a robust expression of Cx26 in control mice (Fig. 1 C and D). We found that recombination in Cx26^fl/fl^ mice significantly reduced the expression of Cx26 induced by injury in the ependyma (Fig. 1 E-G, p<0.0001, Mann-Whitney U test). The number of Cx26 puncta outside the ependyma was similar in control and Cx26^−/−^ mice at 5 DPI (Fig. S3 A-C and E; p=0.45, Kruskal-Wallis multiple comparisons test), showing that the deletion of Cx26 was selective for ependymal cells. We next used a proliferation assay by injecting the thymidine analog EdU from day 2 to 5 after SCI (Fig. 1 B) to test whether the deletion of Cx26 in ependymal cells affected the proliferative reaction of ependymal cells to injury. We found that the number of EdU+ nuclei within the ependyma at 5 days post-injury (DPI) -when injury-induced proliferation peaks-was significantly reduced in the ependyma nearby the lesion epicenter of Cx26^−/−^ mice compared to FoxJ1CreER-tdT control animals (Fig. 1 H-L, ± 250 µm around the lesion epicenter; p<0.0001, Mann-Whitney U test and Fig. S3 F). However, numerous EdU+ nuclei were still observed outside the ependyma (Fig. S3 G-H). Cx26 deletion did not affect the expression of Cx43 in ependymal cells (Fig. S3 J-N).

**Figure 1.**
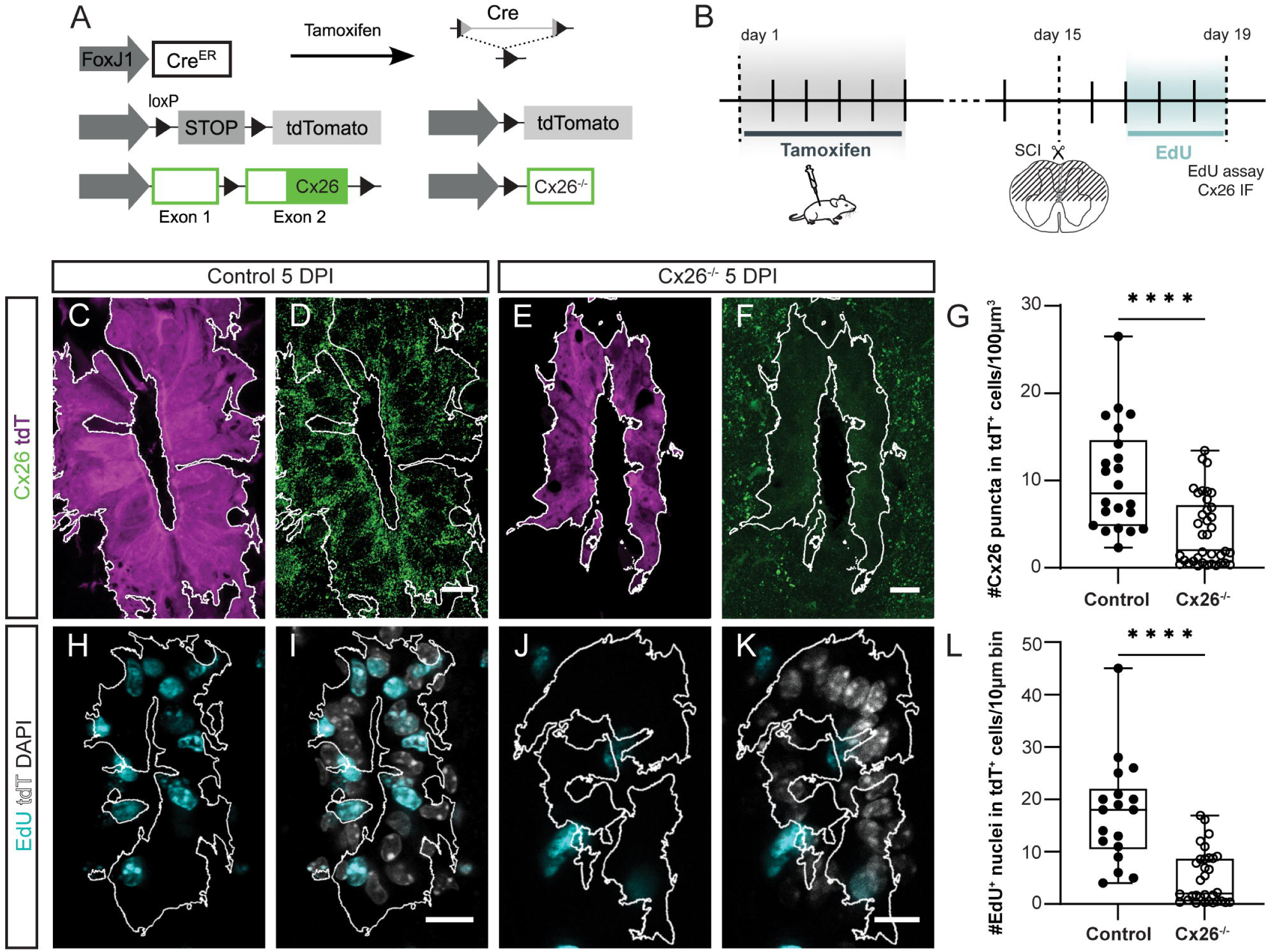
Deletion of Cx26 impairs the injury-induced proliferation of ependymal cells. **A**, genetic strategy to specifically delete Cx26 in adult ependymal cells. **B**, experimental design to analyze the deletion of Cx26 and its impact on the proliferation induced by injury. **C** and **D**, expression of tdT and Cx26 in a FoxJ1CreER-tdT control mouse at 5 DPI, respectively. In this and subsequent figures, the white line delimits the area of tdT expression. **E** and **F**, the expression of Cx26 in the ependyma at 5 DPI is strongly reduced in Cx26^−/−^ mice. **G**, box plot showing a significant difference in the number of Cx26 puncta between control and Cx26^−/−^ mice (p<0.0001, Mann-Whitney U test, n= 6 mice for control and 7 mice for Cx26^−/−^). **H**, EdU+ nuclei in a control mouse at 5 DPI. **I,** merge of EdU+ nuclei and DAPI. **J**, EdU uptake at 5 DPI in a Cx26^−/−^ mouse. **K**, merge of EdU+ nuclei and DAPI. **L**, box plot showing a significant difference in the number of EdU+ nuclei between control and Cx26^−/−^ mice (p<0.0001, Mann-Whitney U test, n= 3 mice for control and 7 mice for Cx26^−/−^). In **H** to **K** the white lines show the mask defined by tdT expression where EdU+ nuclei were counted. Calibration bars: 10 µm.

Various neurotransmitters regulate the biology of ependymal cells (17–20). In particular, ATP is massively released after SCI (21) and the activation of P2X7 receptors (P2X7r) in ependymal cells leads to the resumption of proliferation in a way similar to that induced by injury (22). To explore whether Cx26 is involved in the reactivation of dormant ependymal cells induced by P2X7r activation, we injected the selective P2X7r agonist BzATP (1 mM, 1 µl) close to the central canal (CC) (Fig. S4 A). We compared the proliferation induced by P2X7r activation between FoxJ1CreER-tdT control and Cx26^−/−^ mice. The rate of proliferation in the intact adult ependyma is very low (3, 4, 23) but increases substantially after activation of P2X7r by BzATP injection (22). Figure S4 B-D shows that BzATP injection close to the CC in control FoxJ1CreER-tdT mice induced ependymal cell proliferation as evidenced by EdU uptake. However, genetic deletion of Cx26 strongly reduced the reactivation of ependymal cell proliferation induced by P2X7r activation (Fig. S4 E-G). Figure S4 H shows the spatial profile of EdU uptake around BzATP injection. The analysis of the number of EdU+ nuclei in ± 250 µm around the injection site shows a significant difference in EdU+ nuclei between control and Cx26^−/−^ mice (Fig. S4 I, p<0.0001, Mann-Whitney U test). These data suggest that Cx26 also plays a key role in the molecular mechanisms triggered by activation of P2X7r to induce ependymal cell proliferation.

### Cx43 also contributes to the response of ependymal cells to injury

Unlike Cx26, the expression of Cx43 in ependymal cells close to the injury site does not change after injury (12). Based on wound healing in other tissues (24), we hypothesized that Cx43 might be detrimental for the response of ependymal cells to injury. To evaluate this possibility, we genetically deleted the Cx43 (Gja1) gene from ependymal cells using the same strategy described for Cx26 (Fig. 2 A and B). As for Cx26, Cx43 deletion did not seem to affect the basic characteristics of the ependyma in the uninjured spinal cord (Fig. S2 A, C-D, E, G-H). As previously described (12), Cx43 was robustly expressed in the ependyma of the adult SC of FoxJ1CreER-tdT (Fig. 2 C and D). Recombination in FoxJ1CreER-tdT-Cx43^fl/fl^ mice significantly reduced the number of Cx43 puncta within the ependyma (Fig. 2 E-G, p<0.01 Mann-Whitney U test). Surprisingly, we found that Cx43 deletion reduced the proliferative reaction of the ependyma at 5 DPI (Fig. 2 H-L, p<0.05 Mann-Whitney U test). We wondered whether Cx26 was still being up-regulated by injury when Cx43 had been deleted, but was no longer efficient to promote ependymal cell proliferation. Immunohistochemistry for Cx26 in Cx43^−/−^ mice at 5 DPI showed that in the absence of Cx43, injury did not trigger the Cx26 expression needed to reactivate ependymal cells (Fig. 2 M-Q, p<0.05 Mann-Whitney U test). The expression of Cx26 in the parenchyma outside the ependyma was not affected in Cx43^−/−^ mice (Fig. S3 D and E, p=0.91, Kruskal-Wallis multiple comparisons test). The most parsimonious interpretation of these results is that the effect of Cx43 on injury-induced proliferation is indirect via the interference of the injury-induced Cx26 up-regulation.

**Figure 2.**
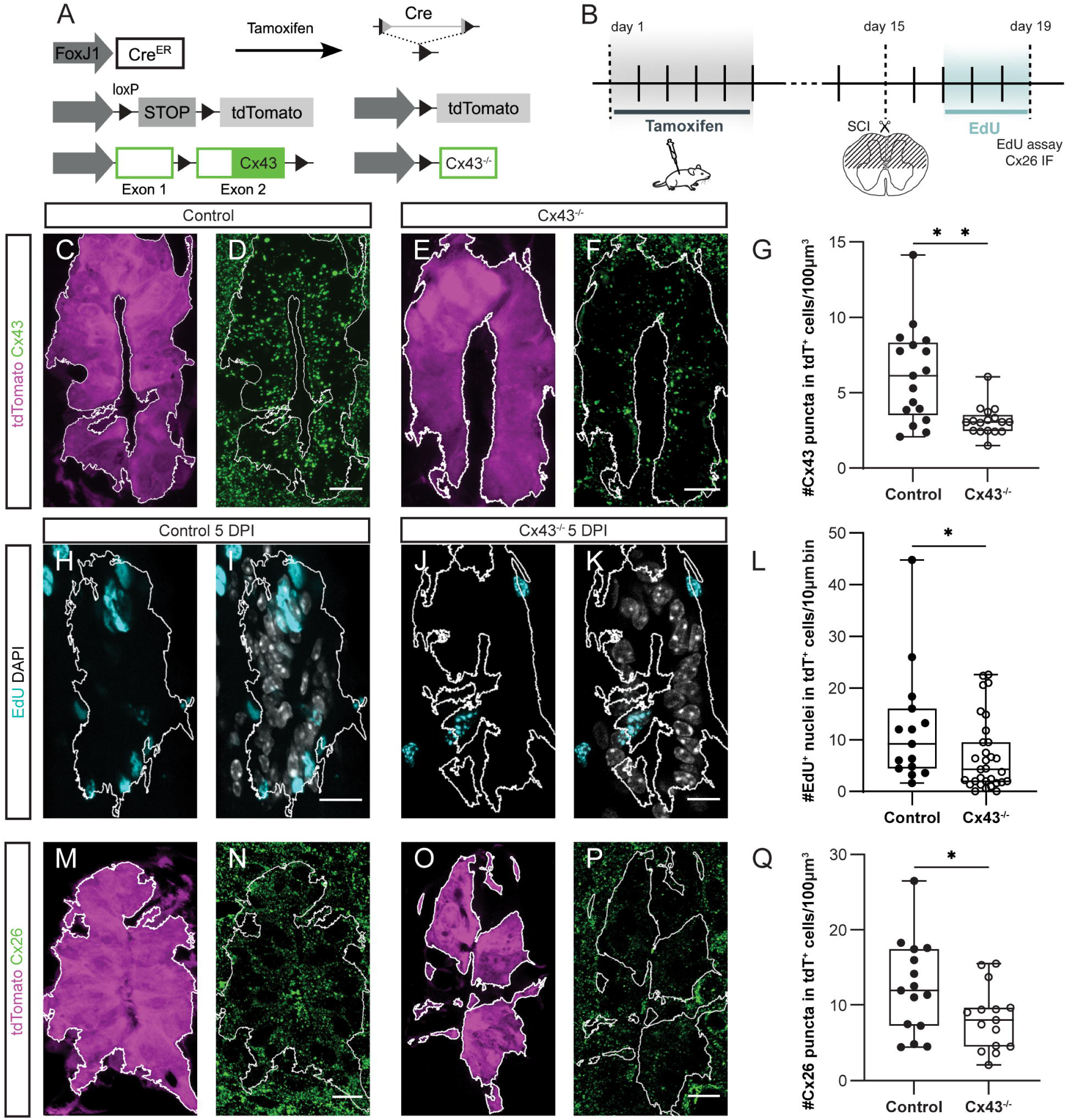
Cx43 also plays a role in the awakening of ependymal cells. **A**, genetic strategy to specifically delete Cx43 in adult ependymal cells. **B**, experimental design to analyze the deletion of Cx43 and its impact on the proliferation induced by injury. **C** and **D**, expression of the reporter gene tdT and Cx43 in a FoxJ1CreER-tdT control mouse, respectively. **E** and **F**, expression of the reporter gene tdT and Cx43 in a Cx43^−/−^ mouse, respectively. **G**, box plot showing that the number of Cx43 puncta in the tdT+ cells is significantly different between control and Cx43^−/−^ mice (p<0.01, Mann-Whitney U test, n=5 mice for each group). **H**, EdU+ nuclei at 5 DPI in a control mouse. **I**, DAPI stained nuclei are overlayed with the EdU+ nuclei shown in **H**. **J**, EdU+ nuclei at 5 DPI in a Cx43^−/−^ mouse. **K**, merge of DAPI and EdU+ nuclei shown in **J**. **L**, box plot showing a statistically significant difference (p<0.05, Mann-Whitney U test, n= 3 mice for control and 5 mice Cx43^−/−^) in cell proliferation between the FoxJ1CreER-tdT control and Cx43^−/−^ mice. **M** and **N**, tdT expression and immunohistochemistry for Cx26 at 5 DPI in a control mouse, respectively. **O** and **P**, tdT and Cx26 expression at 5 DPI in a Cx43^−/−^ mouse, respectively. **Q**, box plot showing there is a significant difference (p<0.05, Mann-Whitney U test, n= 5 mice for control and 4 mice for Cx43^−/−^) in the number of Cx26 puncta between control and Cx43^−/−^ mice at 5 DPI. Calibration bars: 10 µm.

### Communication between ependymal cells by Cxs

We were puzzled by the fact that Cx43 affects Cx26 expression and thereby the response of ependymal cells to SCI. We speculated that the impact of Cx43 deletion on the ependymal cell reaction may be by interference with the communication between ependymal cells. To explore the role of specific Cx isoforms in gap junction coupling among ependymal cells, we exploited the fact that under normal conditions the ependyma of neonatal mice is an active stem cell niche with robust gap junction coupling and expression of both Cx26 and Cx43, allowing the production of numerous spinal cord slices for patch-clamp recordings. As already described for adults, we induced recombination of FoxJ1CreER-tdT-Cx26^fl/fl^ and FoxJ1CreER-tdT-Cx43^fl/fl^ in neonatal mice (two daily injections of tamoxifen at P9 and 10) to delete a selected Cx from ependymal cells (Fig. S5). Patch clamp recordings of neonatal ependymal cells (P11) in control (FoxJ1CreER-tdT) animals revealed cells with passive properties (Fig. S6 A and B, control, n= 12) whose input resistance increased after addition of the gap junction and Cx hemichannel blocker meclofenamic acid (MFA, 100 µM, n= 7; Fig. S6 A-C; p<0.05 Wilcoxon Signed-Rank test), suggesting gap junction coupling and/or functional Cx hemichannels in ependymal cells. In line with the communication of ependymal cells via gap junctions, the diffusion of biocytin -a compound highly permeable through connexons-showed extensive dye coupling (Fig. S6 D). Deletion of Cx26 did not significantly change the input resistance of patch clamp recorded ependymal cells (Fig. S6 G, p=0.97, Mann-Whitney Wilcoxon rank test) or the size of biocytin-labelled cell clusters (Fig. S6 E, H; p> 0.99, Kruskal-Wallis multiple comparison test). In contrast, patch clamp recordings of Cx43^−/−^ ependymal cells showed an increase in the apparent input resistance of cells (Fig. S6 G, p<0.005, Mann-Whitney Wilcoxon rank test) and revealed either isolated cells (Fig. S6 F) or pauci-cellular clusters of coupled cells with significantly smaller volumes (Fig. S6 H; p<0.05, Kruskal-Wallis test). To test whether communication via Cxs in the adult ependyma after injury is similar to that of the neonatal ependyma, we made patch clamp recordings around the lesion epicenter (± 2000 µm; Fig. S6 I). Because both proliferation and Cx26 expression decrease sharply away from the lesion epicenter (12), we performed a multi-linear regression analysis of the input resistance, volume of dye coupled cells and distance from the lesion epicenter that showed a significant effect of genotype on cluster size (F(2,39)=3.589, p=0.037). Compared with the FoxJ1CreER-tdT genotype, Cx43^−/−^ mice displayed a significantly smaller cluster size (p=0.0213, Mann-Whitney test; Fig. S6 J). These data suggests that both in the neonatal as well as in the adult reactive ependyma, Cx43 is the main contributor to communication between ependymal cells via gap junctions.

Besides connexons forming gap junction plaques, there is a sub-population of permeable Cx43 hemichannels in the intact ependyma of the SC (12). In line with this, the uptake of ethidium bromide in ependymal cells of Cx43^−/−^ mice was strongly reduced compared to control FoxJ1CreER-tdT mice (Fig. S7 A-E, p<0.005, Mann-Whitney U test) and in recombined versus non-recombined cells in Cx43^−/−^ mice (Fig. S7 C-D and F, p<0.05, Wilcoxon Signed-Rank test). We then asked whether the effect of Cx43 deletion on the injury-induced proliferation could be partly accounted for by the absence of Cx43 hemichannels. To explore this possibility, we made a pre-emptive blockade of Cx43 hemichannels by means of intraspinal injection of the antagonist peptide TAT-Gap19 (1 mM, 1 µl) before a dorsal hemisection of the cord. The injury site was immediately filled with Pluronic F127 gel embedded with TAT-Gap19 (1 mM, Fig. 3 A). The application of the scrambled version of TAT-Gap19 peptide (Scr-TAT-Gap19) was used as control. We found that Scr-TAT-Gap19 did not affect the injury-induced proliferation (Fig. 3 B and C) but TAT-Gap19 reduced the number of EdU+ nuclei in ependymal cells in response to injury (Fig. 3 D and E). Figure 3 F shows the spatial profile of EdU+ nuclei around the lesion epicenter in TAT-Gap19 and Src-TAT-Gap19. The analysis of EdU+ nuclei in a ± 250 µm extension around the lesion epicenter shows that TAT-Gap19 significantly reduced the injury-induced proliferation of ependymal cells (Fig. 3 G; p<0.0001 Mann-Whitney U test), suggesting that Cx43 hemichannels are an important component of the early signaling required to reactivate the latent ependymal progenitor-like cells.

**Figure 3.**
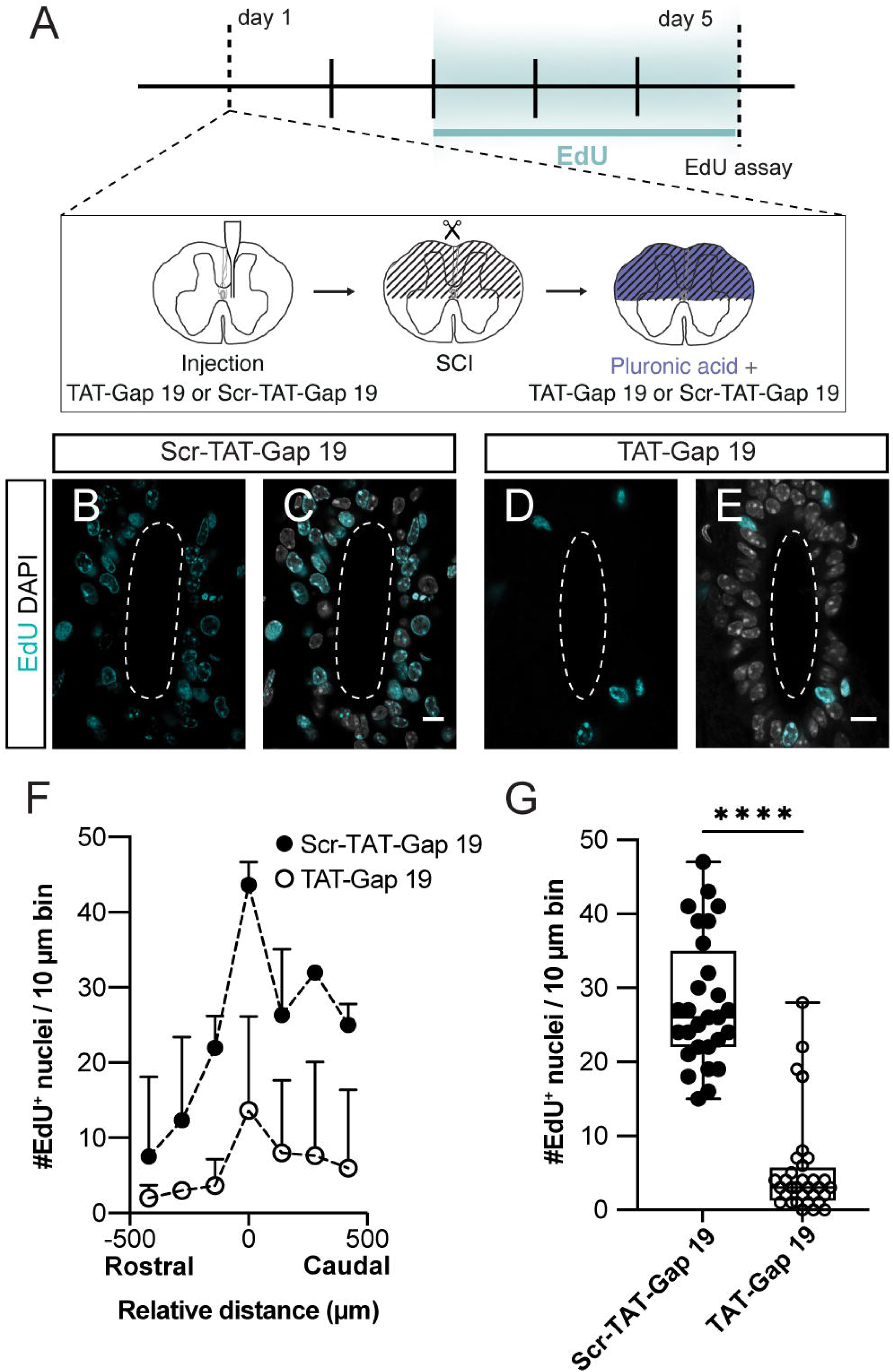
Blockade of Cx hemichannels impaired the reaction of ependymal cells to SCI. **A**, experimental design to explore the role of Cx hemichannels in injury-induced proliferation of ependymal cells. **B**, uptake of EdU in a mouse injected with the scrambled version of TAT-Gap19 at 5 DPI. **C**, DAPI staining overlayed with EdU+ nuclei show a substantial number of proliferating ependymal cells in the presence of Scr-TAT-Gap19. **D**, blockade of Cx hemichannels with TAT-Gap19 strongly reduced the proliferative reaction to SCI. **E**, merge of DAPI and EdU+ nuclei shown in **D**. **F**, spatial profile of EdU+ nuclei around the lesion epicenter. **G**, box plot showing that at 5 DPI the number of EdU+ nuclei in animals treated with TAT-Gap19 is significantly smaller than those treated with Scr-TAT-Gap19 (p<0.0001, unpaired t-test). Calibration bars: 10 µm.

Because Cx43 is expressed in the ependymal cells of the uninjured SC, we speculated that Cx43 may play a part in the earliest events taking place after injury by regulating communication between ependymal cells and Ca^2+^ signals triggered by P2X7r activation (26). To evaluate this possibility, we crossed FoxJ1CreER and Gt(ROSA)26Sortm38(CAG-GCaMP3)Hze/J mice to express GCaMP3 -a Ca^2+^ sensing protein-in ependymal cells (Fig. 4 A) and made spinal cord slices from adult animals (Fig. 4 B). Puff application of BzATP in a discrete region of the CC (Fig. 4 C1) induced Ca^2+^ waves that propagated within the ependyma (Fig. 4 C2-4 and D). In some regions of interest (ROIs), secondary waves appeared at longer delays (Fig. 4 D, asterisks). The secondary waves probably arise from the intrinsic Ca^2+^ dynamics within individual cells as they did not appear to correlate with Ca^2+^ waves in neighboring ROIs. Figure 4 E shows at a faster time scale that the Ca^2+^ wave initiated and peaked earlier in some ROIs, suggesting a propagation from leading to follower cells within the ependyma (Video S1). MFA spared the earliest component but strongly reduced the delayed peak of the Ca^2+^ wave (Fig. 4 F and G). The propagation of the wave measured as the number of ROIs in which ΔF/F_0_ crossed a threshold (F_0_ mean + 8 SD before BzATP) decreased significantly after addition of MFA (100 µM; Fig. 4 H, p<0.01, Wilcoxon Signed-Rank test; Video S2). The comparison of the Ca^2+^ wave propagation among ependymal cells in slices incubated in normal or MFA-containing Ringer’s solution showed a similar result (Fig. 4 I, p<0.005, Mann-Whitney U test) and an overall decrease of the maximal ΔF/F_0_ of different ROIs (Fig. 4 J, p<0.0001 Mann-Whitney U test). Because MFA blocks both gap junctions and Cx hemichannels, we explored the effects of the hemichannel blocker TAT-Gap19 on Ca^2+^ signals triggered by P2X7r activation. Although the mean ratio of responsive ROIs/total ROIs was lower in TAT-Gap19 compared to the scrambled version of the peptide, there was no statistically significant difference (Fig. 4 K, p= 0.5009, Mann-Whitney U test). However, TAT-Gap19 significantly reduced the peak ΔF/F_0_ in the total ROIs analyzed (Fig. 4 L, p<0.0001, Mann-Whitney U test). In line with these data, Ca^2+^ waves monitored with Fluo-4 in ependymal cells (Fig. S8 A-B, C) propagated less (Fig. S8 D and E, p< 0.0005, Mann-Whitney U test, 27 slices for control and 46 slices for Cx43^−/−^ mice) and were smaller in maximal intensity (Fig. S8 D-F, p<0.0001, Mann-Whitney U test, 27 slices for control and 46 slices for Cx43^−/−^ mice) in Cx43^−/−^ mice compared to control animals.

**Figure 4.**
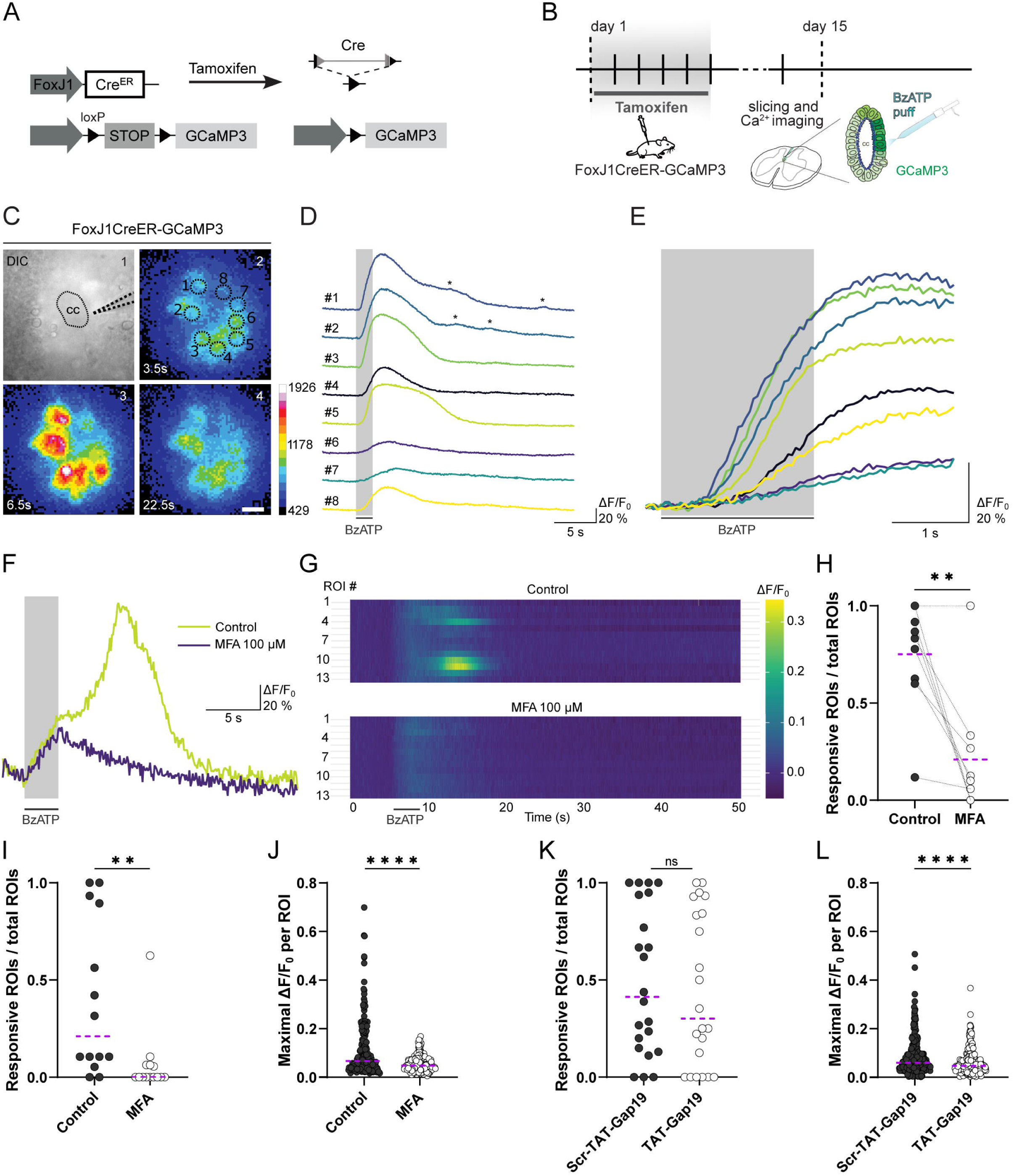
Cx43 hemichannels and gap junctions are needed for Ca^2+^ wave propagation within the ependyma. **A**, genetic strategy to specifically express the Ca^2+^ sensor GCaMP3 in adult ependymal cells. **B**, schematics showing the experimental design to study the Ca^2+^ responses to BzATP application in ependymal cells expressing GCaMP3. **C**, differential interference contrast (DIC) image of an adult SC slice showing the ependyma and the position of the puff pipette containing BzATP (1). Pseudo color images of cells expressing GCaMP3 at different time points (2–4). Regions of interest (ROIs) used for ΔF/F_0_ are indicated with numbered circles. The scale bar shows the fluorescent values in arbitrary units. **D**, fluorescence (ΔF/F_0_) corresponding to each of the ROI shown in **C**. The time of puff application of BzATP is indicated by a black bar. Notice that in some locations secondary peaks in fluorescence appeared (asterisks). **E**, the same data shown in **D** but at a faster time-scale to show the onset of the fluorescence increase, which is dramatically different in different ROIs, suggesting that the calcium signal may travel as a wave within the ependyma. **F**, the late component of the calcium wave was eliminated by the gap junction and hemichannel blocker meclofenamic acid (MFA, 100 µM). **G**, heat map in 13 adjacent ROI showing the dynamic response of the ependyma to BzATP puff application under control conditions and after addition of MFA (100 µM) to the bath. **H**, paired dot plot of the ratio of cells that responded with a Ca^2+^ increase of the total number or cells (responsive ROIs/total ROIs) shows that MFA (100 µM) reduced significantly the propagation of the Ca^2+^ wave within the ependyma (p<0.005, Wilcoxon Signed-Rank test). **I**, ratio of cells responding to BzATP in slices incubated in normal Ringer’s solution (control) or MFA (100 µM) shows a significant difference in Ca^2+^ wave propagation (p<0.005, Mann-Whitney U test, 14 slices per group). **J**, maximum ΔF/F_0_ values for individual ROI shows the incubation in MFA (100 µM) strongly reduced the intensity of Ca^2+^ signals (p<0.0001 Mann-Whitney U test, 14 slices per group). **K**, ratio of cells responding to BzATP in slices incubated in Scr-TAT-Gap19 (200 µM) or TAT-Gap19 (200 µM) shows no significant difference in Ca^2+^ wave propagation (p= 0.5009, Mann-Whitney U test). **L**, maximum ΔF/F_0_ values for individual ROIs in Scr-TAT-Gap19 and TAT-Gap19 incubated slices shows a statistically significant reduction after the blockade of Cx hemichannels (p<0.0001, Mann-Whitney U test, 22 slices per group). Calibration bar: 10 µm.

Although we cannot exclude non-canonical functions of Cx43, our data suggests that Cx43 - probably via gap junctions and hemichannels-is important for the communication of Ca^2+^ signals initiated by activation of P2X7r, blocking downstream events such as the expression of Cx26 that seems critical for the resumption of ependymal cell proliferation.

### Deletion of Cxs in ependymal cells impacts healing

We next evaluated the impact of Cx deletion in ependymal cells on spinal cord repair. We found that the lack of Cx26 up-regulation that would normally occur after injury, interfered with the injury-induced proliferation and had a profound effect on the formation of the glial scar. After Cx26 deletion, fewer tdT+ cells could be observed away from the CC at 15 DPI when compared to control (Fig. 5 A and B). Similar findings were obtained after deletion of Cx43 (Fig. 5 C). The cross-sectional area covered by tdT+ cells over the total area around the lesion epicenter (± 250 µm) was significantly lower in both Cx26^−/−^ and Cx43^−/−^ mice compared to control (Fig. 5 D, p<0.0004 and p<0.0001 respectively, Dunn’s multiple comparisons test). At 15 DPI the lack of Cx26 in ependymal cells impaired the contribution of the ependyma to healing as evidenced by the measurement of the normalized cross-sectional area of the cord around the lesion epicenter compared to the neighboring uninjured cord (Fig. 5 E and F; p<0.05, One Way ANOVA Dunnett’s multiple comparison test). Although the mean normalized cross-sectional area of the cord around the lesion epicenter in Cx43^−/−^ mice was smaller, there was no significant difference compared to control animals (Fig. 5 F, p= 0.53, One Way ANOVA Dunnett’s multiple comparison test). We then explored the effect of Cx26 deletion on healing at 30 DPI when the contribution of the ependyma reaches a plateau (4). In control FoxJ1CreER-tdT mice, the lesion site was filled with scar tissue and abundant ependyma-derived cells at 30 DPI (Fig. 5 G). Many tdT+ cells can be observed migrating away from the CC (Fig. 5 H). The area occupied by tdT+ cells over the total area of the cord at the lesion site remains about the same between 15- and 30-DPI, suggesting that the number of ependyma-derived cells correlates with the restoration of the damaged area (Fig. 5 I). The normalized cross-sectional area around the lesion epicenter was still significantly smaller in Cx26^−/−^ mice compared to control (Fig. 5 J and K, p<0.05, paired t test). Our results are similar to those obtained in mice after deletion of Ras genes in ependymal cells (25), and confirm the importance of the latent ependymal stem cell niche to limit the extension of the lesion and the relevance of Cx signaling to support the ependymal cell response to injury.

**Figure 5.**
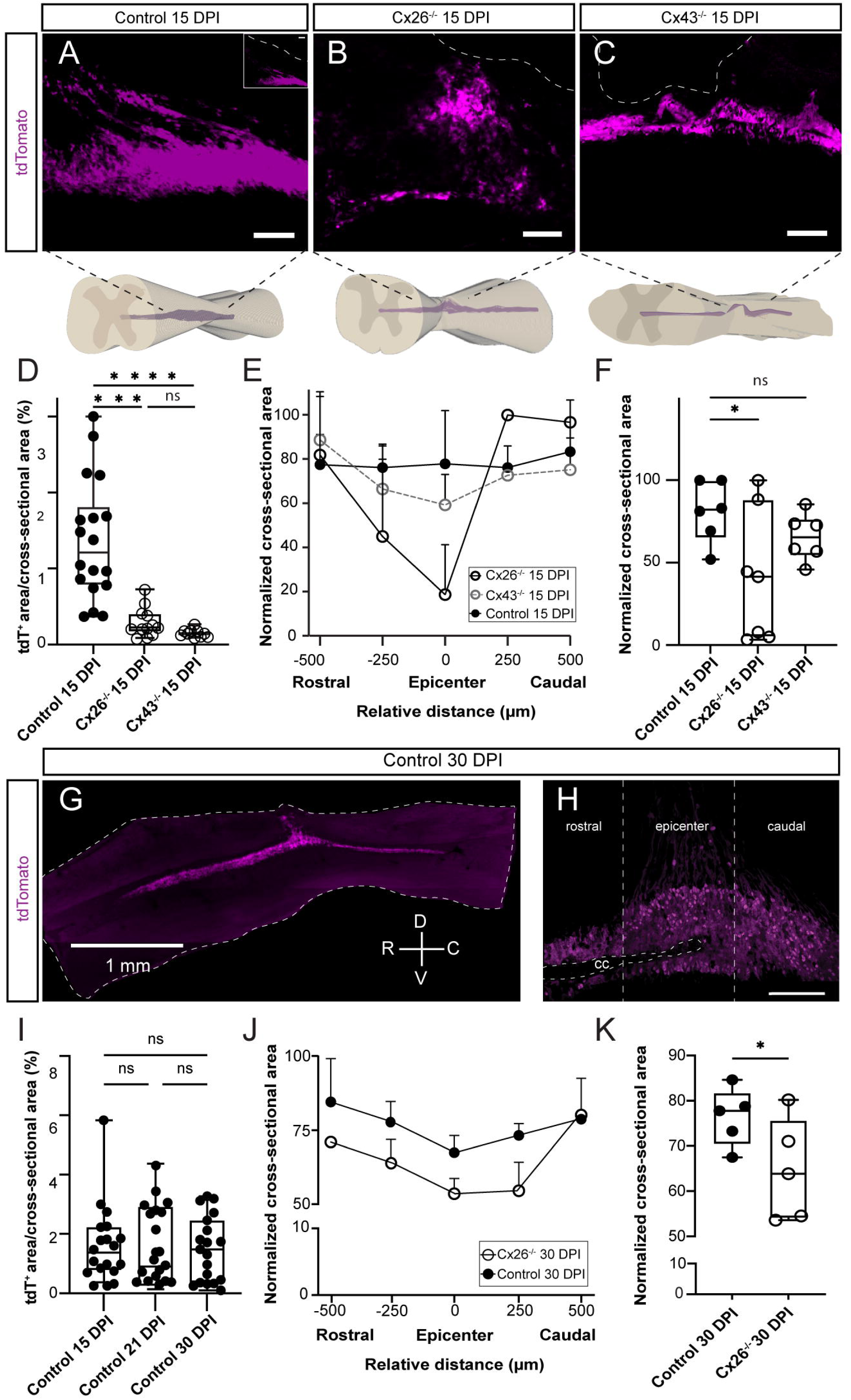
The lack of Cx26 in ependymal cells affects healing. **A** and **B**, representative images of sagittal sections of tdT+ cells around the lesion epicenter in control and Cx26^−/−^ mice at 15 DPI, respectively. **C**, tdT+ cells around the lesion epicenter of a Cx43^−/−^ mouse. The white dotted lines in **A** to **C** show the dorsal limit of the cord whereas the corresponding 3D reconstructions are shown below each panel. **D**, box plot showing the ratio between the cross-sectional area covered by tdT+ cells and the total cross-sectional area in control, Cx26^−/−^ and Cx43^−/−^ mice. **E**, normalized cross-sectional area of the SC around the lesion epicenter (0) in control (FoxJ1CreER-tdT), Cx26^−/−^ and Cx43**^−/−^** mice at 15 DPI. **F**, box plot of the normalized area of the SC around the lesion epicenter (± 250 µm) shows Cx26^−/−^ mice have a significantly smaller cross-sectional area of the cord (p<0.05, One Way-ANOVA Dunnett’s test) whereas there was no significant difference between control and Cx43^−/−^ mice (p=0.53, One Way-ANOVA Dunnett’s test, n= 3 mice for each group). **G**, sagittal section of the spinal cord of a FoxJ1CreER-tdT mouse at 30 DPI. The dotted line highlights the outer limit of the cord. **H**, tdT+ cells of a FoxJ1CreER-tdT mouse migrating away from the CC around the lesion epicenter (vertical dotted lines). **I**, box plot of the tdT+ area over the total cross-sectional area of the cord of FoxJ1CreER-tdT mice at 15, 21 and 30 DPI. There were no significant differences between selected time points (15 vs. 21 DPI, p> 0.99; 15 vs. 30 DPI, p> 0.99; 21 vs 30 DPI, p>0.99; Kruskal-Wallis Multiple Comparisons test; n= 3 mice for each group). **J**, normalized cross-sectional area of the SC around the lesion epicenter (0) in control (FoxJ1CreER-tdT) and Cx26^−/−^ mice at 30 DPI. **K**, box plot of the normalized cross-sectional area of the cord around the lesion epicenter in control and Cx26^−/−^ mice at 30 DPI (p<0.05, paired t test). Calibration bars: **A-C** and **H**, 100 µm; **G**, 1000 µm.

## Discussion

Endogenous latent progenitor-like cells in the ependyma are a potential source for improving healing and functional recovery after SCI (5). To optimize the potential of the ependymal stem cell niche for repair, understanding the mechanisms by which progenitors are reactivated by injury is fundamental. Cxs play key roles in the regulation of the biology of stem cells and in wound healing in different tissues. Here, we show that Cx26 and Cx43 are needed for the reaction of ependymal cells to injury, and their lack has profound implications for endogenous repair.

Different isoforms of Cxs are dynamically expressed from the earliest embryonic developmental stages and regulate events such as proliferation, migration and differentiation of neural progenitors (27). In particular, Cx43 and Cx26 exhibit specific spatiotemporal expression patterns in the mouse cortical plate, regulating radial glia proliferation and the migration of newborn neurons (7,8). The importance of Cxs is maintained in adult neurogenic niches as knockouts of Cx30 and Cx43 in progenitors of the dentate gyrus decrease their proliferation (9). Both Cx43 and Cx26 are expressed in ependymo-radial glia of the spinal cord of the turtle, a vertebrate with remarkable regenerative capabilities (10, 28). Similarly, the active ependymal stem cell niche of neonatal mice has a robust expression of Cx26 and Cx43 with a prominent electrical and metabolic communication via gap junctions, suggesting that Cx signaling in the ependyma shares mechanisms with bona fide stem cell niches of the developing mammalian brain (8, 29, 30). As postnatal development proceeds and the ependyma becomes quiescent, gap junction coupling and Cx26 expression decline, whereas Cx43 remains unchanged. Gap junction coupling and Cx26 expression increases around the lesion epicenter at the time injury-induced proliferation peaks, suggesting a role for Cx26 in the resumption of the mitotic activity of ependymal cells (12). In line with this, we find here that selective genetic deletion of Cx26 in ependymal cells dramatically reduced the proliferative reaction of the ependymal stem cell niche to injury. The reduced number of ependyma-derived cells had a profound effect on scar formation. Our results resemble those of Sabelström et al. (2013) (25) obtained by deletion of Ras genes and support the idea that the contribution of the ependymal stem cell niche is fundamental to limit the extension of tissue damage. Cx26 and Cx43 are differentially expressed during the cell cycle of murine neocortical precursors, with Cx26 increasing in mitotic cells between the S and early G1 phases (7, 8). Since Cx43 expression in ependymal cells around the lesion epicenter remains unchanged after injury, our results suggest that a low Cx26/Cx43 ratio in adult ependymal cells characterizes a non-proliferative progenitor phenotype in the intact cord, that shifts to a proliferative state with the rise of Cx26 after injury. Similar to injury, the activation of proliferation induced by P2X7r is paralleled by the re-expression of Cx26 (22). The fact that genetic deletion of Cx26 blocked the effect of P2X7r activation on the induction of proliferation suggests that Cx26 is a fundamental hub for the signaling pathways needed to keep progenitor-like cells of the ependyma in an active state.

Cx26 plays a key role in the development and wound healing of different tissues. For example, selective Cx26 ablation arrests the postnatal development of the organ of Corti (31) and impairs the development of excitatory neocortical neurons (32). In the normal skin, Cx26 expression is very low but it is upregulated around the wound between 1 to 4 days after skin injury (14). Likewise, Cx26 is associated with proliferation in healing of human airway epithelial cell cultures (33). Similarly, our findings support the idea that Cx26 is a key regulator of the proliferation of progenitor-like cells in the ependyma in the context of tissue repair after SCI. Besides proliferation, Cxs have been shown to play a role in cell migration (34) and recently Cx43 has been implied in the migration of ependyma-derived cells (19). The expression of Cx26 in HeLa cells increases cell motility in scrape wounding models by reducing cell adhesion (35). It may be possible that Cx26 in ependymal cells is involved in the detachment and migration of ependyma-derived cells. However, the strong interference of Cx26 deletion on injury-induced proliferation precluded the study of an eventual role in the migration of ependyma-derived cells. The molecular mechanisms by which Cx26 is upregulated and exert its modulation on proliferation of ependymal cells remain to be explored. Interestingly, during pregnancy Cx26 is strongly upregulated both in mammary epithelial and uterine endometrial cells by transcriptional regulation via SP and AP-2 transcription factors (36, 37). The present study suggests that Cx26 is not the main Cx isoform mediating gap junction coupling in the ependyma. This raises the possibility that the effect of Cx26 on proliferation is mediated by non-canonical functions via the interaction with other proteins. Interestingly, in breast cancer cells Cx26 forms a complex with NANOG and focal adhesion kinase to drive self-renewal (38). Future studies should explore potential protein interactions with Cx26 to understand the molecular mechanisms by which the cell cycle of ependymal cells is regulated.

The functional role of Cx43 in the healing of different types of epithelia is complex as it regulates cellular events such as migration and proliferation (39). The persistence of Cx43 under pathological conditions impairs healing (40) whereas blockade of Cx43 improves wound closure in the skin (41) and superficial corneal epithelial wounds (42). In the SC, Cx43 is strongly expressed in astrocytes and play a role in the spread of tissue damage as its blockade promotes some functional recovery (43, 44). However, our loss-of-function approach showed that instead of an improvement, Cx43 deletion in ependymal cells decreased the reaction of the ependymal stem cell niche to injury. The fact that Cx43 deletion reduced the response of ependymal cells to injury at 5 DPI but did not have a significant impact on healing at longer survival times, suggests that beyond early signaling of tissue damage, Cx43 may have other functions that influence repair as reported for some epithelia (40–42).

Although we cannot rule out the interaction of Cx43 or its carboxyterminal with other proteins (45, 46), our data suggests that the effect of Cx43 deletion on the reactivation of the latent ependymal stem cell niche by injury can be accounted for by the interference of Cx26 upregulation. To fully resolve this issue, future studies should explore whether forcing the expression of Cx26 in ependymal cells can rescue the Cx43^−/−^ phenotype. Electrophysiology and dye coupling indicate that Cx43 is the main Cx isoform responsible for the communication between ependymal cells. In addition, in the intact spinal cord there is a subpopulation of functional Cx43 hemichannels (12) and P2X7r (26). Our data shows that P2X7r activation by BzATP induced Ca^2+^ waves that spread within the ependyma, a propagation that depended on Cx signaling as it was blocked by MFA. The differential effects induced by MFA and TAT-Gap19 suggests that Cx43 hemichannels may be involved in the generation of full Ca^2+^ signals whereas gap junctions may affect primarily the propagation of waves. As in many other cell types, Cx43 hemichannels may release ATP that would then activate P2X7r in an auto- and paracrine way. The Ca^2+^ influx through P2X7r would induce a Ca^2+^-induced Ca^2+^ release from the endoplasmic reticulum via ryanodine receptors (26) and permeation of Ca^2+^ through gap junctions, leading to the spread of Ca^2+^ among a cohort of ependymal cells. The increase in intracellular Ca^2+^ would then activate the transcription of Cx26 to reactivate dormant progenitors in the ependyma.

Collectively, our findings show that Cx26 represents an essential hub for the signaling pathways that lead to the resumption of proliferation within the ependymal stem cell niche in response to injury. We propose the main role of Cx43 is -together with P2X7r activation by ATP- to initiate a sequence of cellular events by facilitating the spread of Ca^2+^ signals, leading to Cx26 upregulation in a cohort of ependymal cells. Cx26 appears as a relevant target to manipulate the ependyma towards a functional state more competent to promote endogenous repair. Because ependyma-derived cells do not produce axon inhibiting compounds (e.g., chondroitin sulphate proteoglycans, ref. 4) but generate neurotrophic factors that promote neuronal survival after injury (25), it is tempting to speculate that the overexpression of Cx26 may increase the contribution of the ependyma to repair, thereby limiting the extension of the damage and leading to a more permissive scar. A gain-of-function approach should be used in future studies to test this possibility.

## Material and Methods

### Animals

FoxJ1CreER-R26RtdTomato mice (kind gift of Prof. Jonas Frisén, Karolinska Institutet) (47) were used as control. B6.129S7-Gja1^tm1Dlg^/J (The Jackson Laboratories, RRID:IMSR_JAX:008039) and B6;129P2-Gjb2^tm1Ugds^/Cnrm (The European Mouse Mutant Archive - EMMA, EM:00245) mice were crossed with FoxJ1CreER-R26RtdTomato (FoxJ1CreER-tdT) mice to selectively delete Cx43 or Cx26 from ependymal cells, respectively. B6.Cg-Gt(ROSA)26Sor^tm38(CAG-GCaMP3)Hze^/J (The Jackson Laboratories, RRID:IMSR_JAX:029043) mice were crossed with FoxJ1CreER to express the fluorescent calcium indicator protein GCaMP3 selectively in ependymal cells. To induce Cre dependent recombination in adult mice (P>90), we injected tamoxifen (Sigma; 2 mg, 20 mg/ml in corn oil, i.p.) for 5 days and allowed at least 10 days between the last injection and surgery to ensure clearance (47). For neonatal mice, two injections of tamoxifen were applied at P9-10 and processed at P11. Male and female animals were used indistinctly. All experimental procedures were approved by our local Committee for Animal Care (protocol #005/09/2015).

### Spinal cord injury (SCI)

Animals were anesthetized using isoflurane (3% for induction, 1-1,5% for maintenance, Terrel^TM^). Injury of the dorsal aspect of the SC was performed as described elsewhere (4,12). Briefly, after laminectomy the dorsal aspect of the spinal cord at a low thoracic level (T13) was cut transversely with microsurgical scissors deepened 0.8 mm into the cord to achieve a dorsal hemisection that included the ependyma at the lesion epicenter. The lesion was extended rostrally for about one spinal cord segment by means of longitudinal cuts along the midline. To minimize variability all lesions were performed by the same researcher (M.V.F.). Tramadol (3 mg/kg, i.p) was administered for pain relief immediately after surgery. A second dose of tramadol was applied 24 h after surgery.

### *In vivo* activation of P2X7 receptors (P2X7r)

Intraspinal injection of the selective P2X7r agonist BzATP (1 μl, 1 mM in saline) was performed as described elsewhere (22). Briefly, the injection was performed with a glass micropipette (∼40 μm tip) pulled from graduated glass capillaries (GC-3.5, RWD Life Sciences). The glass micropipette was secured in a microinjector (RWD life Sciences, R-480 Nanoliter Microinjection Pump) attached to a stereotaxic holder. The pipette was lowered 550 μm lateral from the midline to a depth of 700 μm to be placed nearby the ependyma without inducing a mechanical damage. The drug was injected at a speed of 0.1 μl/min and the retention time was set in 10 min. After the injection, the pipette was withdrawn in 100 μm steps at 15 s intervals.

### *In vivo* blockade of Cx43 hemichannels

Intraspinal injection of the selective Cx43 hemichannel blocker TAT-Gap19 (YGRKKRRQRRR-KQIEIKKFK; 1 μl, 1 mM in saline, GenScript) or the scrambled peptide scr-TAT-Gap19 (GEIRRIRYKKKFKQKRRQKR; 1 µl, 1 mM in saline, GenScript) was performed as described for the in vivo activation of P2X7 receptors and elsewhere (22). After the removal of the micropipette, we performed a dorsal hemisection which was immediately followed by the injection within the lesion site of TAT-Gap19 (1 mM) or ScrTAT-Gap19 (1 mM) dissolved in Pluronic F-127 (20% in PBS, 50 µl).

### Proliferation assay

A proliferation assay was done as previously described (12, 22). Briefly, EdU (40 mg/kg, i.p.; Biosynth Ltd.) was injected twice daily (4 h interval) from day 3 to day 5 after BzATP injection or SCI. Animals were sedated with diazepam (10 mg/kg, i.p.) and anesthetized with ketamine (100 mg/kg, i.p) and xylazine (10 mg/kg, i.p.) to be perfused with 4% paraformaldehyde in 0.1 M phosphate buffer (PB, pH 7.4). EdU labelling was revealed by using the Click-iT Alexa Fluor 647 imaging kit (Invitrogen, Thermo Fisher Scientific).

### Slice preparation and electrophysiology

Neonatal or adult mice were anesthetized with isoflurane (Terrel^TM^). Immediately after achieving full unresponsiveness to painful stimuli, mice were decapitated and the low thoracic-upper lumbar spinal cord dissected out under cutting Ringer’s solution. For neonatal animals the cutting Ringer’s solution had the following composition (in mM): NaCl, 101; KCl, 3.8; MgCl_2_, 18.7; CaCl_2_, 1; MgSO_4_, 1.3; HEPES, 10; KH_2_PO_4_, 1.2 and glucose, 25; saturated with 5% CO_2_ and 95% O_2_ (pH 7.4). Transverse 300 µm slices of the thoraco-lumbar spinal cord were made. To analyze the properties of the adult reactive ependymal cells, the injured spinal cord at 5 DPI was isolated and a transverse cut with microscissors was performed at the level of the lesion epicenter. Then, both the rostral and caudal stumps were placed facing upwards and transverse spinal cord slices (200-300 µm) were cut in a Ringer’s solution with the following composition (in mM): NMDG, 92; KCl, 2.5, NaH_2_PO_4_, 1.5; NaHCO_3_, 30; HEPES, 20; Glucose, 25, thiourea, 2; Na-Ascorbate, 5; Na-Pyruvate, 3 CaCl_2_, 0.5, MgSO_4_, 10; saturated with 5% CO_2_ and 95% O_2_, pH 7.4). Slices were incubated for 15 min at 34°C in the same cutting Ringer’s solution before storage in resting Ringer’s solution (in mM, NaCl, 119; KCl, 2.5, NaH_2_PO_4_, 1.5; NaHCO_3_, 26; glucose, 12.5, thiourea, 2; Na-Ascorbate, 5; Na-Pyruvate, 3 CaCl_2_, 2, MgSO_4_, 10, saturated with 5% CO_2_ and 95% O_2_ to keep pH 7.4) at room temperature until their use. Slices were placed in numbered wells of a storage chamber in order to correlate their position with respect to the lesion epicenter. For patch clamp recordings, slices were placed in a recording chamber and superfused at 1 ml min^−1^ with normal extracellular Ringer’s solution (in mM: NaCl, 124; KCl, 2.4; NaHCO_3_, 26; CaCl_2_, 2.4; MgSO_4_.6H_2_O, 1.3; HEPES, 1.25; KH_2_PO_4_, 1.2; and glucose, 10; saturated with 5% CO_2_ and 95% O_2_, pH 7.4). All recordings were performed at room temperature (22°-24°C). Cells were visualized with differential interference contrast optics (Leica Microsystems, DM LFS) or oblique illumination (Olympus, BX51WI) and patch-clamp whole-cell recordings obtained with electrodes filled with (in mM): K-gluconate, 122; Na_2_-ATP, 5; MgCl_2_, 2.5; CaCl2_2_, 0.003; EGTA, 1; Mg-gluconate, 5.6; K-HEPES, 5; H-HEPES, 5; and biocytin, 10; pH 7.3, 5–10 MΩ). Voltage-clamp recordings were performed with a Multiclamp 700B amplifier using pClamp10 (Molecular Devices) or a HEKA amplifier (EPC10 USB) using Patchmaster. Seal resistances were between 1 and 4 GΩ. In voltage-clamp mode, cells were held at −70 mV and the resting membrane potential (RMP) was estimated from the current-voltage relationship (at I= 0). The input resistance was calculated around the RMP.

### Cx hemichannel permeability assay

The permeability of Cx hemichannels in ependymal cells was evaluated by the uptake of the hemichannel permeable dye ethidium bromide (EtBr; 4 µM). Briefly, spinal cord slices from Cx43^−/−^ and control mice were obtained as described above and incubated in Ringer’s solution devoid of Ca^2+^ 15 min before and during the application of EtBr (15 min). After incubation, slices were rinsed in normal Ringer’s solution for 5 min, then fixed overnight in 4% PFA in PB (pH 7.4), rinsed in PB (3 times, 10 min) and finally mounted in glycerol for imaging. Ependymal cells were imaged in an LSM800 Zeiss Airyscan confocal microscope. Stacks of consecutive confocal images were taken at 1 µm intervals with the 561 nm laser for tdT (detection wavelengths of 545–605 nm), the laser 640 nm for methyl green (detection wavelengths of 600–700 nm), and the laser 405 nm for EtBr (detection wavelengths of 450–700 nm). Dye uptake analysis was performed in a region of interest defined by the outline of the ependyma. EtBr uptake (as fluorescence units, FU) was expressed as the difference between the fluorescence measured in ependymal cells and the background fluorescence from a field with no labeled cells. Nuclei were counter-stained with methyl green (4 µg/ml, 1 h). For internal control, non-recombined ependymal cells were identified as tdT-, methyl green+ (48) within the ependymal cell layer.

### Morphological identification of patch-clamp recorded cells

During whole-cell patch-clamp recordings, cells were filled with biocytin. After completion of the recording period (15 to 30 min), slices were fixed by immersion in 4% PFA in 0.1M phosphate buffer (PB, pH 7.4) overnight. Following PB rinsing, the slices were incubated in PB containing 0.3% Triton X-100 with streptavidin-Alexa488 or −546 (1:200; Invitrogen, Thermo Fisher Scientific), mounted in Prolong (Invitrogen) and imaged in a confocal microscope (Carl Zeiss, LSM800 Airyscan).

### Ca^2+^ imaging

To visualize calcium dynamics, FoxJ1CreER mice were crossed with 6.Cg-Gt(ROSA)26Sortm38(CAG-GcaMP3)Hze/J mice (JAX #014538). Recombination was induced with tamoxifen, and imaging performed 14 days later. Slices from adult animals (300 µm thickness) were obtained as described above and separated into 4 groups that contained either normal Ringer’s solution (control) or Ringer’s solution containing meclofenamic acid (MFA, 100 µM), TAT-Gap19 (200 µM) or TAT-Gap19 scrambled peptide (200 µM). In some experiments, the Ca^2+^ indicator Fluo4-AM was used to monitor Ca^2+^ signals. Briefly, spinal cord slices from control or Cx43^−/−^ mice were incubated (60 min) in normal Ringer’s solution containing 5 µM Fluo4-AM (dissolved in 20% Pluronic F-127 in DMSO). After wash out of Fluo4-AM (30-60 min) the slices were placed in a recording chamber for imaging. Ca^2+^ transients were evoked by focal puff application of BzATP (500 µM, 1-3 s duration). Time-lapse imaging (5-20 Hz, 430-510 nm excitation filter) was performed using either a LucaR or an iXon EMCCD (Andor) camera using Imaging Workbench 6.0 (INDEC BioSystems) or SOLIS 4.31 (Oxford Instruments) software, respectively. Regions of interest (ROIs) were manually drawn around the central canal in areas exhibiting GCaMP3 or Fluo-4 fluorescence. For each ROl, the background-corrected fluorescence F was calculated as the mean pixel intensity (mean pixel value ROI (t) - mean pixel value ROI background (t)). Baseline fluorescence F_0_ was defined as the average F within 3 seconds before the BzATP application. ΔF/F_0_ was calculated for each ROI as (F−F_0_)/F_0_. In Fluo-4 recordings, linear correction was applied to the ΔF/F_0_ data using the baseline interval as reference in order to improve signal analysis. For event (peak) detection, a responsive ROI was defined as that displaying an event exceeding a threshold. In the case of GCaMP3 recordings, the threshold was calculated for each ROI individually (F_0_ + 8SD (F_0_)) whereas for Fluo-4 a fixed threshold of ΔF/F_0_= 0.10 was applied uniformly across all ROIs. To evaluate the propagation of the Ca^2+^ wave within the ependyma, the ratio of ROIs crossing the threshold over the total number of ROIs (# responsive ROIs/total # ROIs) was calculated for every slice. The maximum ΔF/F_0_ value for each ROI was also calculated. Analysis of the time-lapse images for ROI determination and measurement was performed using FIJI (49). Background and linear corrections, ΔF/F_0_ calculation and peak detection were performed using custom R scripts (50).

### Immunohistochemistry

Animals were anesthetized with ketamine (100 mg/kg, i.p), xylazine (10 mg/kg, i.p.) and diazepam (10 mg/kg, i.p.) and fixed by intracardiac perfusion with 4% paraformaldehyde in 0.1 M PB. The spinal cord was sectioned with a vibrating microtome (70 μm thick), washed twice in PBS and then incubated with primary antibodies in PBS with 0.3% Triton X-100 (Sigma-Aldrich). The following primary antibodies were used: anti-Cx43 (rabbit polyclonal, 1:500; Alomone Labs, #ACC-201); anti-Cx26 (rabbit polyclonal, 1:200; Alomone Labs, #ACC-212), Ki67 (rabbit polyclonal, 1:1000, Abcam, ab15580), Sox2 (goat polyclonal, 1:100; Santa Cruz Biotechnology, sc-17320). Sections were then incubated in secondary antibodies conjugated with different fluorophores. Nuclei were stained with DAPI or Hoechst (Invitrogen). Control experiments were performed omitting primary antibodies.

### Experimental design and statistical analyses

Adult (P90 to P180) or neonatal (P9 to P11) mice of either sex from the strains mentioned above were assigned to experimental groups. Injured mice were included in the study only when paralysis of both hindlimbs resulted immediately after recovery from the anesthesia. Mice were excluded from the study when loss of weight at the time of euthanasia -following the procedures approved by our institutional Committee for Animal Care (protocol #005/09/2015)-was above 15% of the initial weight, when the lesion anatomically exceeded 0.8 mm or was less than 0.7 mm from the dorsal aspect of the cord and when recombination efficiency was lower that 60% measured as the tdT+ area/total ependymal area (76.86 ± 13.13 %, n=35). All images obtained for quantitative analysis were taken using the same pre-set parameters and analyzed with Fiji. For quantitative analysis of immunofluorescence data, a minimum of 5 sections were analyzed and averaged per biological replicate (3–5 animals for each experimental condition, unless otherwise stated). To analyze changes in the expression of Cx26 and Cx43, the immunoreactive puncta were converted into particles using a macro developed in IJ1macro language (51) and quantified in the area occupied by tdT around the lesion site (± 250 μm, in 70 μm transverse sections). The confocal images were acquired with an LSM800 Zeiss Airyscan confocal microscope. To quantify the degree of healing after injury, the thickness of the cord around the lesion site was measured as the normalized cross-sectional area relative to the area of the most proximal intact spinal cord (20). Statistical analyses were performed with GraphPad prism 9.50.0 with statistical significance set at P<0.05. The statistical test chosen for each experiment is noted next to the corresponding result. Numerical values are reported as mean ± SD. SD. Statistically analyzed data have been deposited in a public repository (52).

## Acknowledgments

We would like to thank Dr. Jonas Frisén for the generous gift of the FoxJ1CreER-tdTomato transgenic mouse. We thank Tabaré de los Campos for developing a macro to quantify connexin puncta.

