## Supplementary Figures for "Connexins are essential for the contribution of latent progenitors to healing after spinal cord injury"

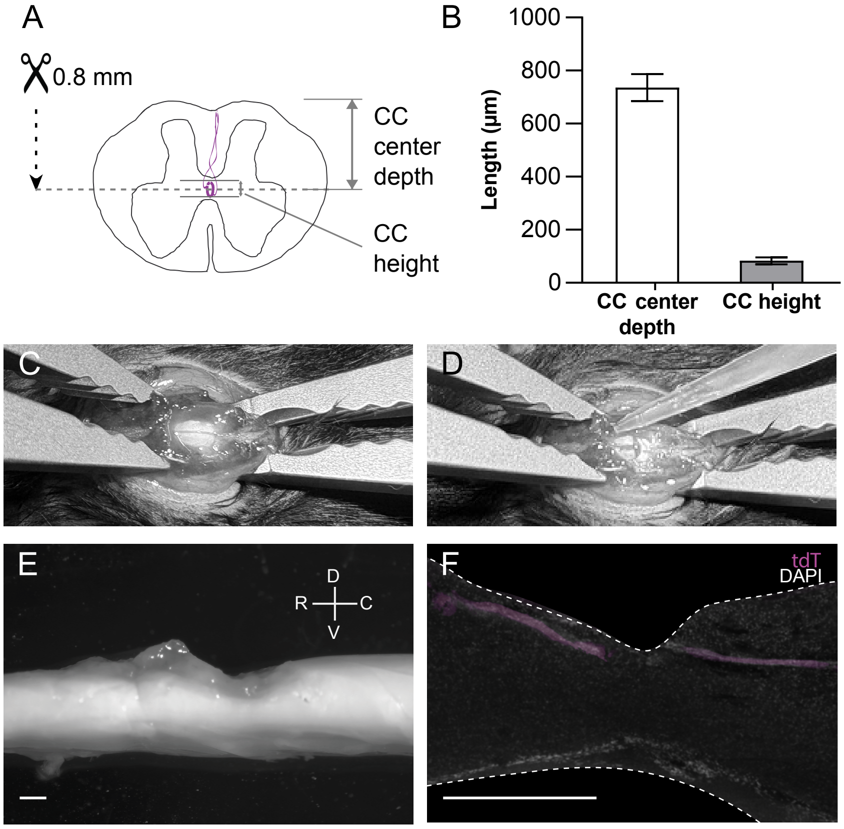


**Supplementary Figure 1. Spinal cord injury model: dorsal hemisection.** **A**, schematic drawing showing that the dorso-ventral extent of the cut (0.8 mm) compromises the ependyma. **B**, bar plot showing the average distance from the dorsal aspect of the cord to the ependyma (735.2 ± 51 µm, n= 18) and the average dorso-ventral length (82.63 ± 13.09 µm, n=18) of the ependyma. **C**, picture showing the dorsal aspect of the spinal cord after a laminectomy. The spinal cord is immobilized by means of two clamps. **D**, same as in **C** after a dorsal hemisection. **E**, panoramic view of the fixed injured spinal cord at 5 DPI. **F**, para-sagittal section of the spinal cord showing the extent of the lesion 24 h after a dorsal hemisection. Notice that the lesion reaches the ependyma (tdT signal) at the epicenter. Calibration bars: **E** and **F**, 1 mm.

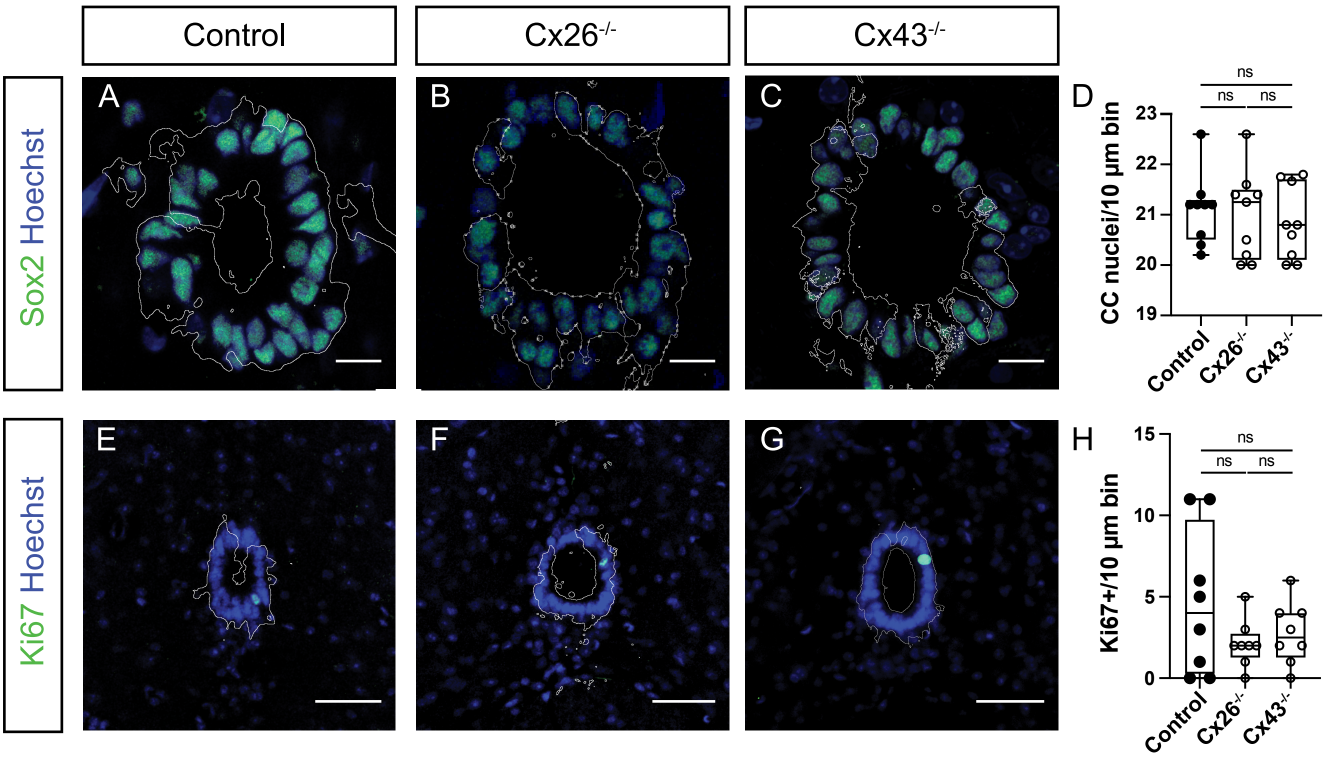


**Supplementary Figure 2. The ependyma of the uninjured spinal cord in control, Cx26$^{-/-}$ and Cx43$^{-/-}$ mice share basic properties. A** to **C**, expression of Sox2 shares the same pattern in the ependyma of control, Cx26$^{-/-}$ and Cx43$^{-/-}$ mice. Nuclei are counterstained with Hoestch. **D**, the number of nuclei in 10 µm bins is similar in control and Cx26$^{-/-}$ (p= 0.48, Mann Whitney U test), control and Cx43$^{-/-}$ (p= 0.32, Mann Whitney U test) and Cx26$^{-/-}$ and Cx43$^{-/-}$ (p= 0.74, Mann Whitney U test) mice. **E** to **G**, immunohistochemistry for Ki67 in the ependyma of control, Cx26$^{-/-}$ and Cx43$^{-/-}$ mice, respectively. **H**, plot showing that there was no difference in Ki67 immunostaining between control, Cx26$^{-/-}$ and Cx43$^{-/-}$ mice.

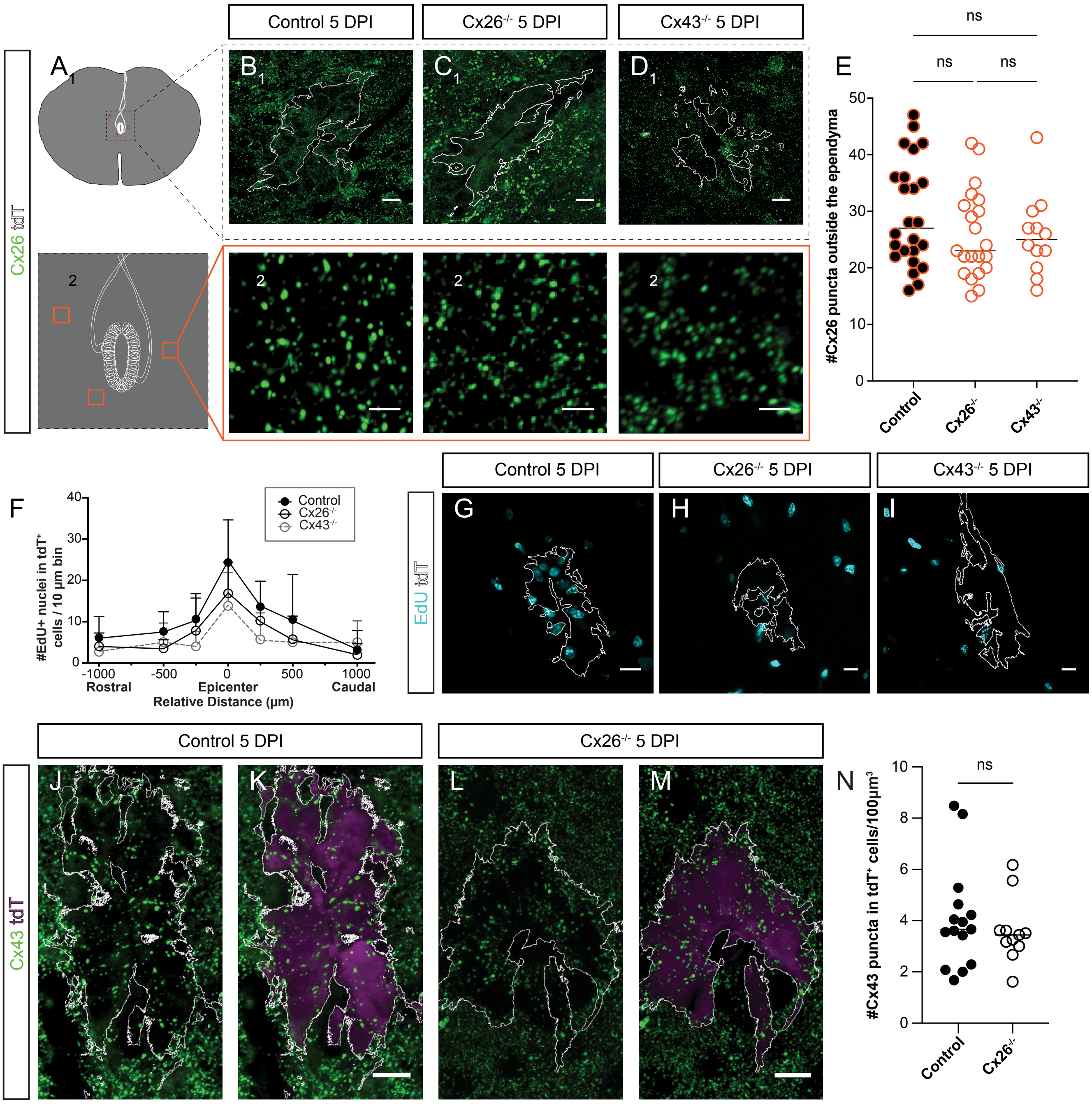


**Supplementary Figure 3. Deletion of Cx26 and Cx43 is specific for ependymal cells. A**, schematic drawing showing the region (dotted line square) around the central canal (1) and the regions outside the ependyma (orange squares in 2) where Cx26 immunoreactivity was sampled. **B-D**, Cx26 immunoreactivity at 5 DPI around the ependyma (1) and far from the ependyma (2), for control, $Cx26^{-/-}$, and $Cx43^{-/-}$ mice, respectively. Notice in **C** the lack of Cx26 puncta within the tdT mask but the abundant puncta outside the ependyma (1). Conspicuous Cx26 puncta are observed far from the ependymal region (2) in all genotypes. **E**, dot plots showing that there were no significant differences in the density of Cx26 puncta outside the ependyma between control and $Cx26^{-/-}$ ($p = 0.45$, Kruskal-Wallis multiple comparison test with post hoc Dunn's test), or control versus $Cx43^{-/-}$ ($p=0.91$, Kruskal-Wallis multiple comparison test with post hoc Dunn's test) and $Cx26^{-/-}$ versus $Cx43^{-/-}$ ($p>0.99$, Kruskal-Wallis multiple comparison test with post hoc Dunn's test) mice. **F**, plot showing the number of EdU+ nuclei per 10 µm around the lesion epicenter (± 1000 µm) in control, $Cx26^{-/-}$ and $Cx43^{-/-}$ mice. **G** to **I**, EdU+ nuclei in the region surrounding the central canal. The white lines define a mask delimiting tdT expression. Notice that while the number of Edu+ nuclei within the mask are less in $Cx26^{-/-}$ and $Cx43^{-/-}$ mice compared to control, there are still abundant EdU+ nuclei outside the ependyma. **J**, Cx43 immunohistochemistry in a FoxJ1CreER-tdT mouse at 5 DPI. **K**, merge of tdT expression in ependymal cells and Cx43 depicted in panel J shows abundant Cx43 puncta within the ependyma. **L**, Cx43 immunohistochemistry in a FoxJCreER-tdT-$Cx26^{-/-}$ mouse at 5 DPI. **M**, merge of tdT expression and Cx43 depicted in panel **L**. Notice the numerous Cx43 puncta in the ependyma of the $Cx26^{-/-}$ mouse. **N**, dot plot showing that the number of Cx43 puncta in control and $Cx26^{-/-}$ mice are not statistically different ($p = 0.413$, Mann-Whitney U test). Calibration bars: **B1, C1** and **D1**, **G** to **I**; **J** to **M**: 10 µm, **B2, C2** and **D2**: 2 µm. The white lines in some panels show the mask defined by tdT expression.

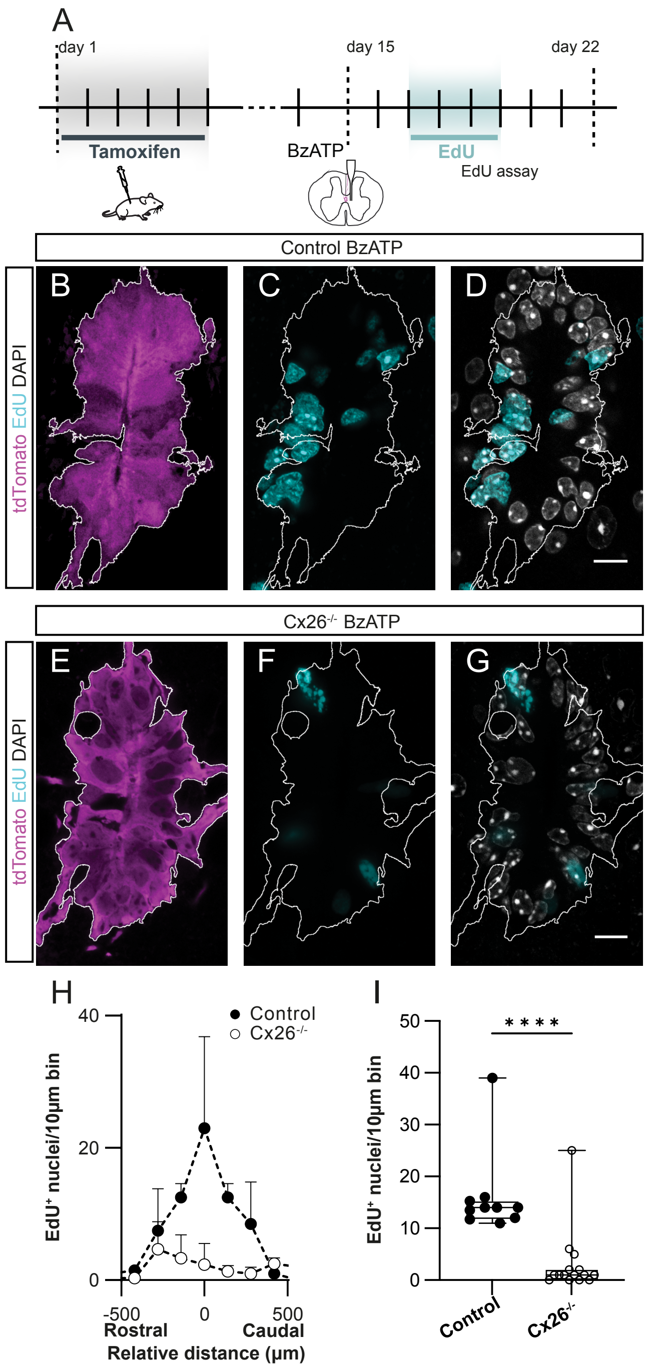


**Supplementary Figure 4. Cx26 is needed for the reactivation of the ependyma induced by purinergic signaling. A**, experimental design to study the role of P2X7r on the activation of quiescent ependymal cells. **B**, recombined ependymal cells expressing tdT in a FoxJ1CreER-tdT control animal. **C**, BzATP injection in control mice induced a strong proliferative response. **D**, merge of DAPI stained and EdU+ nuclei. **E-G**, upon deletion of Cx26 in ependymal cells, BzATP injection failed to reactivate proliferation. **H**, spatial profile of the EdU+ nuclei around the site of BzATP injection. **I**, box plot showing that the number of EdU+ nuclei in BzATP injected animals was significantly higher in control compared to $Cx26^{-/-}$ mice ($p<0.0001$, Mann-Whitney U test, n= 3 mice for each group). Calibration bars: 10 µm.

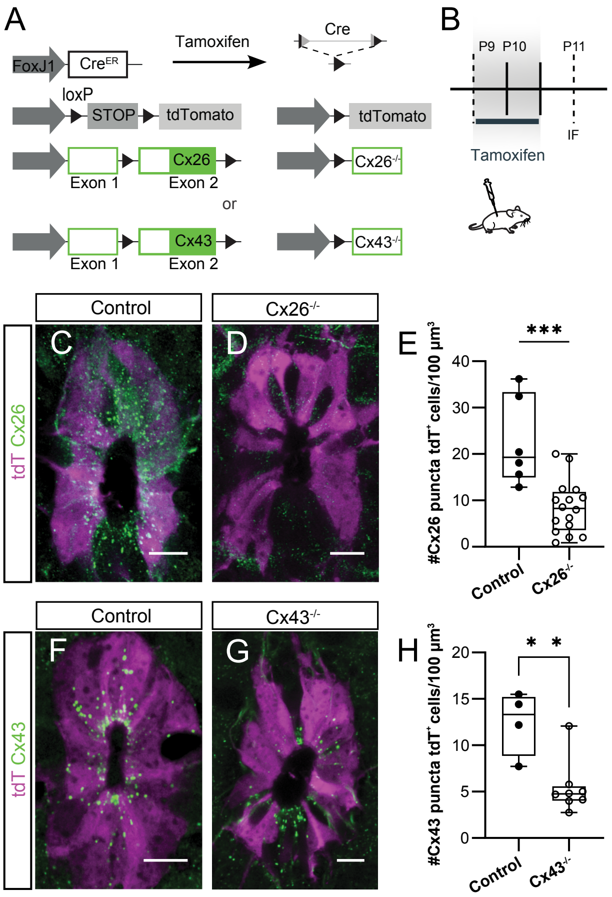


**Supplementary Figure 5. Deletion of Cx26 and Cx43 in the active ependymal stem cell niche of neonatal mice. A**, genetic strategy to specifically delete Cx26 or Cx43 in ependymal cells of neonatal mice. **B**, schematics of the experimental design. **C**, immunohistochemistry for Cx26 in FoxJ1CreER-tdT mice. **D**, expression of Cx26 in the ependyma is strongly reduced in Cx26$^{-/-}$ neonatal mice. **E**, box plot showing a significant difference in the number of Cx26 puncta between control and Cx26$^{-/-}$ mice ($p<0.005$, Mann-Whitney U test). **F**, immunohistochemistry for Cx43 in the ependyma of a control mouse. **G**, in a Cx43$^{-/-}$ mouse the number of Cx43 puncta was strongly reduced. **H**, box plot showing a significant difference in the number of Cx43 puncta between control and Cx43$^{-/-}$ mice ($p<0.01$, unpaired t-test).

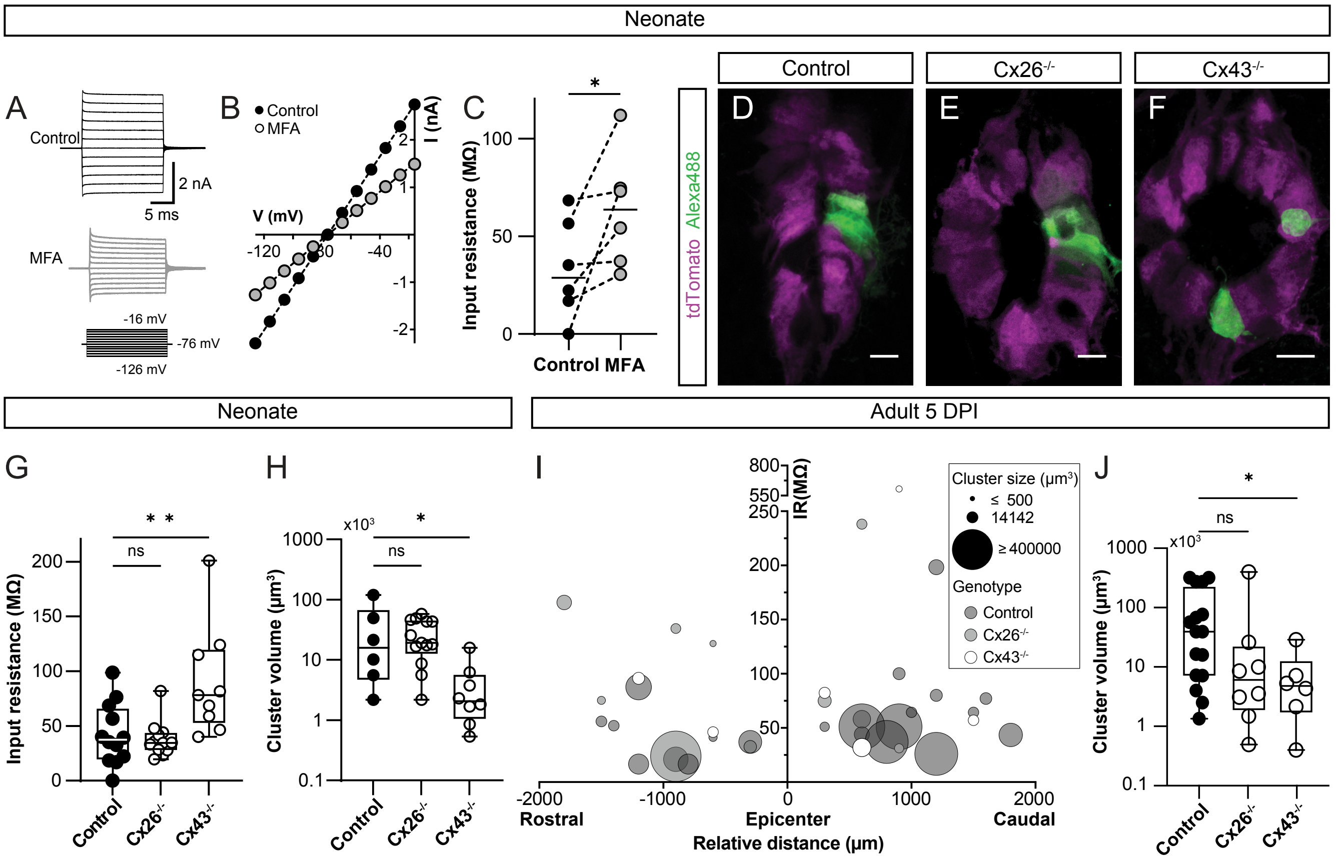


**Supplementary Figure 6. Communication via gap junctions in the intact active neonatal and adult injured ependymal stem cell niches. A**, currents induced by a family of voltage steps in an ependymal cell of a neonatal FoxJ1CreER-tdT mouse in control medium and after addition of meclofenamic acid (MFA, 100 µM). **B**, current-voltage relationship of data shown in **A**. **C**, paired dot plot showing a significant increase in input resistance induced by MFA (100 µM; $p<0.05$ Wilcoxon Signed-Rank test), **D** to **F**, representative images of biocytin filled cells recovered after patch-clamp recordings in control, Cx26$^{-/-}$ and Cx43$^{-/-}$ neonatal mice, respectively. In panel **F** two different patch clamp recordings were performed in the same SC slice. **G**, plots showing there was no significant difference between the input resistance of recorded cells between control and Cx26$^{-/-}$ mice ($p=0.97$, Mann-Whitney Wilcoxon rank test). However, the input resistance of cells in Cx43$^{-/-}$ mice was significantly higher than control ($p<0.005$, Mann-Whitney Wilcoxon rank test). **H**, box plot of the volume of dye coupled cells shows that clusters in Cx43$^{-/-}$ neonatal mice were significantly smaller than control ($p<0.05$, Kruskal-Wallis multiple comparison test). There was no significant difference in the volume of dye coupled ependymal cells between control and Cx26$^{-/-}$ neonatal mice ($p> 0.99$, Kruskal-Wallis multiple comparison tests). **I**, multi-linear regression analysis combining the input resistance, volume of dye coupled cells and distance from the lesion epicenter in adult mice. The plot shows a significant effect of the FoxJ1CreER-tdT-Cx43$^{-/-}$ genotype on cluster size ($F(2,39)=3.589$, $p=0.037$). **J**, the size of dye coupled cell clusters in FoxJ1CreER-tdT-Cx43$^{-/-}$ in ependymal cells of the adult injured spinal cord was significant smaller compared to control mice ($p=0.0213$, Mann-Whitney test). Cluster size in the FoxJ1CreER-tdT-Cx26$^{-/-}$ genotype was not statistically different to that of control mice ($p=0.12$, Mann-Whitney U test). Calibration bars: 10 µm.

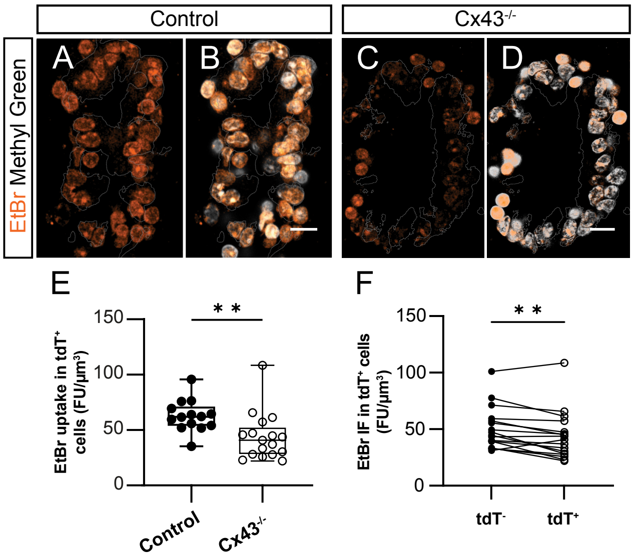


**Supplementary Figure 7. Functional Cx43 hemichannels in ependymal cells of the adult spinal cord. A**, EtBr uptake in control mice (FoxJ1CreER-tdT). **B**, overlay of DAPI stained nuclei and the EtBr signal shown in **A**. **C**, EtBr uptake in a Cx43$^{-/-}$ mouse. **D**, overlay of DAPI stained nuclei and the EtBr signal shown in **C**. **E**, box plot showing the EtBr uptake in Cx43$^{-/-}$ mice was significantly lower than in control mice ($p<0.005$, Mann-Whitney U test, n= 3 for each group). **F**, comparison of the EtBr uptake in recombined (tdT+) and non-recombined (tdT-) ependymal cells of Cx43$^{-/-}$ mice ($p<0.005$, Wilcoxon Signed-Rank test). Calibration bars: 10 µm.

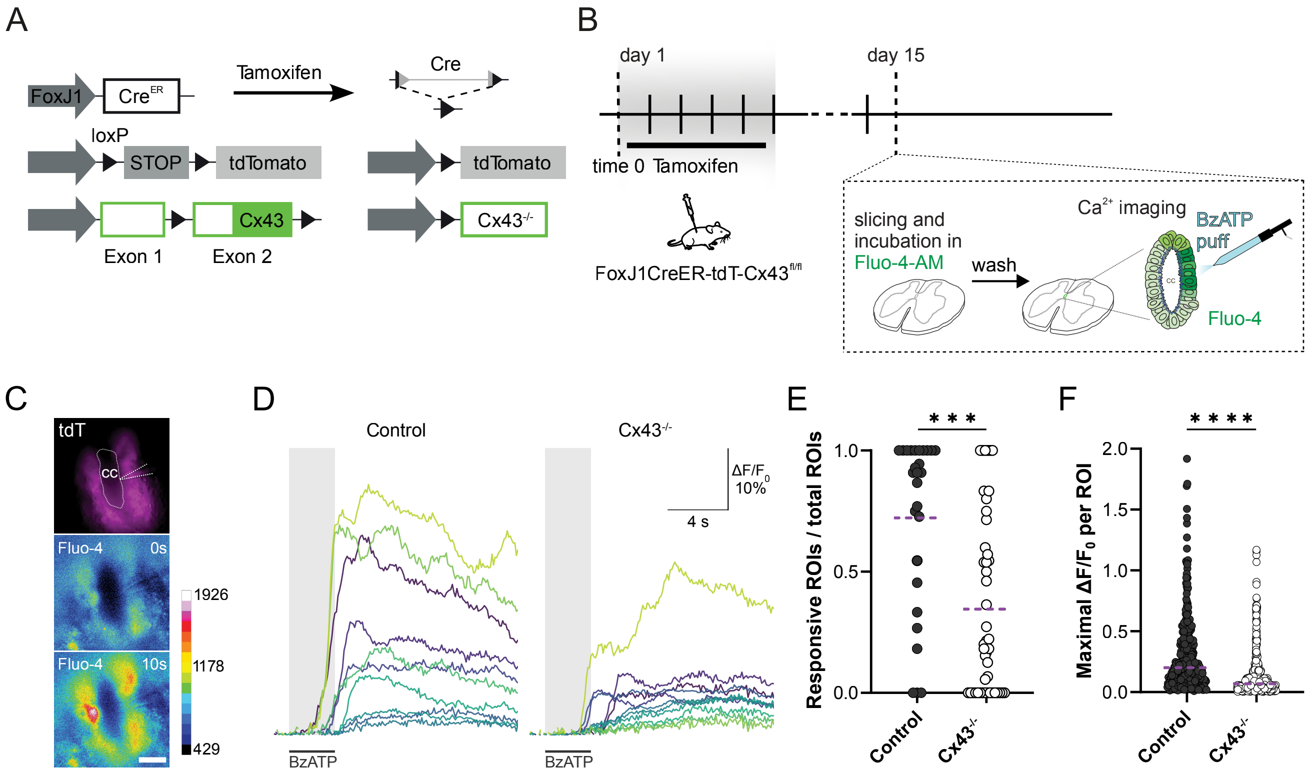


**Supplementary Figure 8. Cx43 deletion in ependymal cells impairs Ca2+ signaling.** **A**, strategy to genetically delete Cx43 in adult ependymal cells. **B**, experimental design to study $Ca^{2+}$ signals in ependymal cells of $Cx43^{-/-}$ mice using the $Ca^{2+}$ sensor Fluo-4. **C**, representative images of the ependyma of a FoxJ1-CreER-tdT mouse (upper panel) loaded with Fluo-4 before and 10s after application of BzATP. **D**, fluorescence ($\Delta F/F_0$) corresponding to different ROIs within the ependyma in control and $Cx43^{-/-}$ mice upon application of BzATP. The time of puff application of BzATP is indicated by a black bar. **E**, the ratio of cells responding to BzATP in slices from control and $Cx43^{-/-}$ mice was significantly different ($p < 0.0005$, Mann-Whitney U test, 27 slices for control and 46 slices for $Cx43^{-/-}$ mice). **F**, the maximum $\Delta F/F_0$ values for individual ROI from control and $Cx43^{-/-}$ mice shows a statistically significant reduction when Cx43 was deleted ($p<0.0001$, Mann-Whitney U test, 27 slices for control and 46 slices for $Cx43^{-/-}$ mice). Calibration bar: 10 μm. In **E** circles represent measurements in individual slices whereas in **F** represent individual ROIs from spinal cord slices shown in **E**. Magenta dotted lines represent the mean.

**Supplementary Video 1**. **$Ca^{2+}$ wave propagation within the adult ependyma**. The DIC image shows the position of the BzATP puff pipette close to ROI #2. The $\Delta F/F_0$ corresponding to different ROIs are displayed on the right in synchrony with the time lapse image. Notice that ROI #3 has an initial rise and a later increase in $Ca^{2+}$ after 10 s followed by a plateau phase. Calibration bar: 10 µm.

**Supplementary Video 2**. **Connexin-signaling modulate $Ca^{2+}$ waves in the ependyma**. $Ca^{2+}$ wave propagation and peak $\Delta F/F_0$ intensity decrease in the presence of the gap junction blocker meclofenamic acid (100 µM). Calibration bar: 10 µm.
